# Structure-activity relationship studies of three novel 4-aminopyridine K^+^ channel blockers

**DOI:** 10.1101/731943

**Authors:** Sofia Rodríguez-Rangel, Alyssa D. Bravin, Karla M. Ramos-Torres, Pedro Brugarolas, Jorge E. Sánchez-Rodríguez

## Abstract

4-Aminopyridine (4AP) is a specific blocker of voltage-gated potassium channels (K_V_1 family) clinically approved for the symptomatic treatment of patients with multiple sclerosis (MS). It has recently been shown that [^18^F]3F4AP, a radiofluorinated analog of 4AP, also binds to K_V_1 channels and can be used as a PET tracer for the detection of demyelinated lesions in rodent models of MS. Here, we investigate three novel 4AP derivatives containing methyl (-CH_3_), methoxy (-OCH_3_) and trifluoromethyl (-CF_3_) in the 3 position as potential candidates for PET imaging and/or therapy. We characterized the physicochemical properties of these compounds (p*K*_a_ and logD) and analyzed their ability to block Shaker K^+^ channel under different voltage and pH conditions. Our results demonstrate that all three derivatives are able to block voltage-gated potassium channels. Specifically, 3-methyl-4-aminopyridine (3Me4AP) was found to be approximately 7-fold more potent than 4AP, whereas the methoxy (3MeO4AP) and trifluoromethyl (3CF_3_4AP) containing compounds were about 3- to 4-fold less potent than 4AP, respectively. These results suggest that these novel derivatives are potential candidates for therapy and imaging.

## Introduction

In normally myelinated neurons, voltage-gated potassium (K^+^) channels K_v_1.1 and K_v_1.2 are clustered near the nodes of Ranvier beneath the myelin sheath^1,2^. Upon demyelination, these channels become exposed, migrate through the demyelinated segment and concomitantly increase in expression several fold^3–9^. This aberrant redistribution of K^+^ channels impairs conduction of action potentials, which leads to neurological deficits^3,10–14^. 4-aminopyridine (4AP) is a selective blocker of K_v_ channels^15–21^used clinically to improve neurological conduction in people with multiple sclerosis (MS)^22–26^ and other demyelinating diseases^27,28^. Mechanistically, 4AP blocks the exposed K^+^ channels and therefore enhances conduction^16,17,19,20,29–32^. Recently, it has been shown that the fluorinated derivative 3-fluoro-4-aminopyridine (3F4AP) also binds to these channels^33^ and, once labeled with ^18^F, can serve to detect areas of demyelination using positron emission tomography (PET)^33–37^. Given the potential of these molecules as therapeutic and imaging agents, we set out to investigate three new 4AP derivatives and their structure-activity relationships.

Prior work on the structure-activity relationship studies of 4AP derivatives has shown that small modifications on the 3 position are permitted^33,38–40^ and that the *in vivo* potency is highly correlated with the p*K*_a_^29,31^. 4AP and derivatives are basic compounds that exist in the protonated or neutral form depending on the pH of the medium (**Fig. 1**). The protonated form mimics a large K^+^ ion and is required to block the channel^41^, while the neutral form is required for the drug to get across the blood-brain barrier (BBB)^42^. Therefore, a p*K*_a_ near the physiological pH is ideal. In addition, the pharmacokinetic properties are largely dependent on the lipophilicity and p*K*_a_ of these molecules. For example, 4AP has a p*K*_a_ of 9.6, resulting in high potency but slow penetration into the CNS, which explains why a slow release formulation is required for therapy^22,23,43^. 3F4AP, on the other hand, has a p*K*_a_ of 7.6 which results in much faster CNS penetration and good favorability for PET imaging^33^. Additional aminopyridine derivative examples include 3,4-diaminopyridine (3,4-DAP), a potent K_v_ channel blocker^40^ with low BBB permeability^42^, used clinically for Lambert-Eaton syndrome^44^, a disorder of peripheral nervous system, and 4-aminopyridine-3-methanol (3MeOH4AP) which has been shown to enhance conduction in laboratory models of spinal cord injury and MS^39,45^ but that has minimal permeability to the BBB and requires intrathecal administration.

**Fig. 1.**
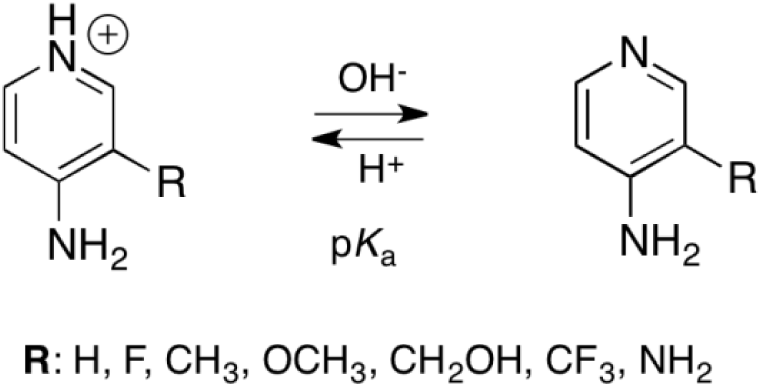
Acid-base equilibrium of 4-aminopyridine derivatives.

Given the correlations between the p*K*_a_, lipophilicity and molecular size with *in vivo* activity, we hypothesized that the derivatives 3-methyl-4-aminopyridine (3Me4AP), 3-methoxy-4-aminopyridine (3MeO4AP) and 3-trifluoromethyl-4-aminopyridine (3CF_3_4AP) would be permeable to the CNS and be suitable candidates for therapy and/or imaging (**Figure 2A**). These molecules are particularly interesting as potential PET radioligands since they are amenable to labeling with ^11^C, which provides some advantages over ^18^F-labeled radioligands. While ^11^C has a significantly shorter half-life compared to ^18^F (20 *vs*. 110 min) limiting its use to sites with a cyclotron, ^11^C-labeled tracers tend to be easier to radiolabel and their short half-life allows for multiple scans on the same subject and day. In fact, we recently communicated methods to produce [^11^C]3MeO4AP and [^11^C]3CF_3_4AP^46,47^ as well as PET imaging results with [^11^C]3MeO4AP showing similar distribution as [^18^F]3F4AP in primate brains^46^.

**Figure 2.**
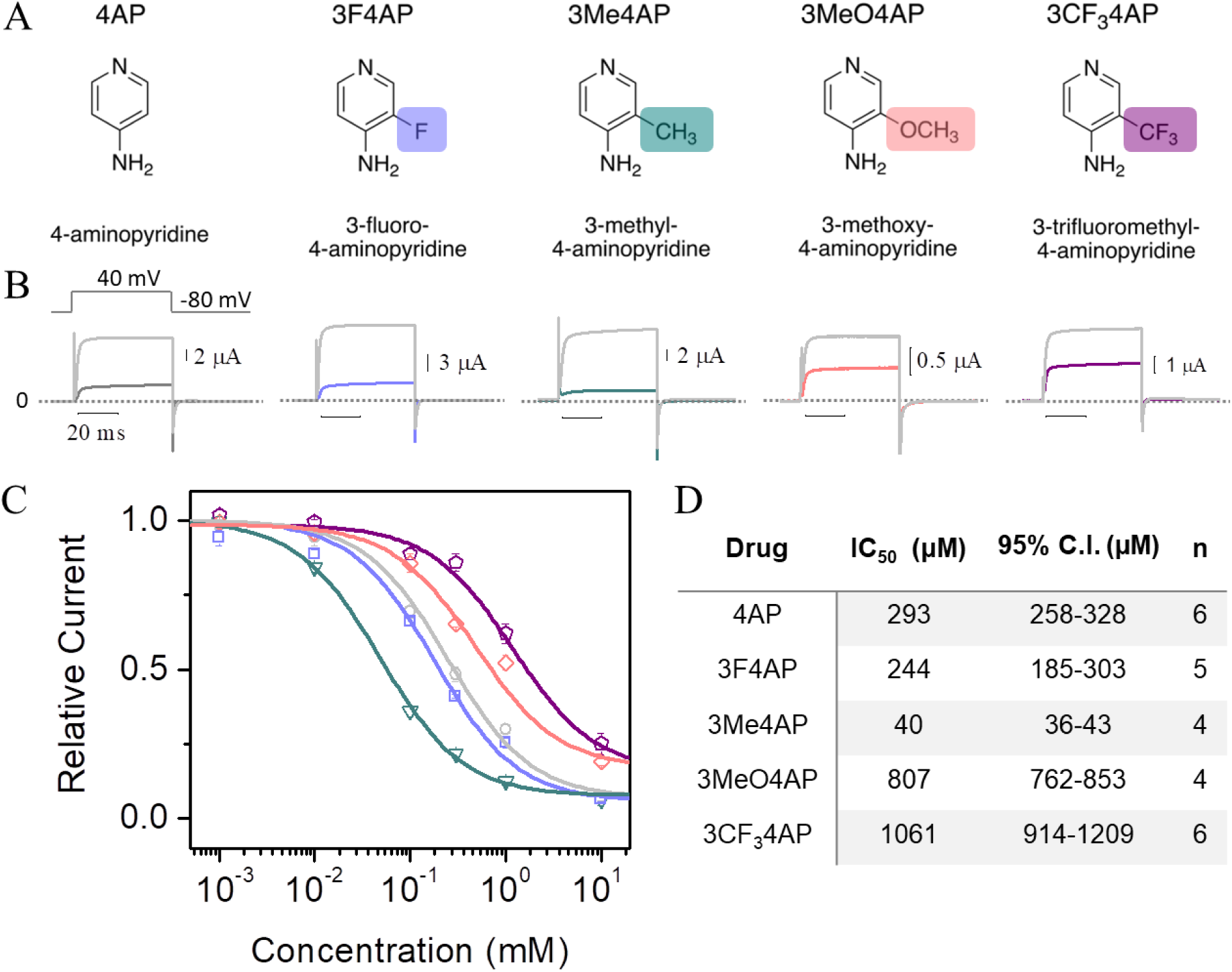
Inhibition of K^+^ currents by 4AP analogs. (**A**), Chemical structures of studied 4AP analogs. From left to right: 4-aminopyridine (4AP), 3-fluoro-4-aminopyridine (3F4AP), 3-methyl-4-aminopyridine (3Me4AP), 3-methoxy-4-aminopyridine (3MeO4AP), 3-trifluoromethyl-4-aminopyridine (3CF_3_4AP). (**B**), Representative recordings of K^+^ current acquired from four to six different oocytes expressing the Shaker K_v_ ion channel before (gray) and after (colored line) addition of 1 mM of each 4AP analog. Currents were elicited by 50 ms depolarization steps from −100 to 50 mV in increments of 10 mV. For clarity, only the inhibition of K^+^ current by cumulative concentration of 4AP and derivatives recorded at 40 mV is shown. The dashed line indicates current at a value of zero and the 20 ms horizontal bar represents the time scale for all recordings. (**C**), Relative current as a function of the concentration of 4AP analogs obtained at 40 mV. (**D)**, IC_50_ of each 4AP analog and 95% confidence interval obtained by fitting the data with the Hill equation (Eqn 1). n represents the number of times each drug was tested in separate oocytes. IC_50_ was determined at pH 7.4.

## Methods

### pK_a_ determination

The p*K*_a_ of each compound was measured by acid titration. Briefly, 5 mg of each compound were dissolved in 5 mL of water and titrated with 0.01 M HCl solution. pH was monitored with a pH meter and plotted as a function of volume of acid added to make a titration curve. p*K*_*a*_ was then found using Gran plot analysis for each replicate. This procedure was repeated 4 times for each compound.

### Partition coefficient determination

The octanol-water partition coefficient (logD) at pH 7.4 was determined using a modified version of the shake flask method. Briefly, PBS (900 µL), 1-octanol (900 µL) and a 10 mg/mL aqueous solution of each compound (2 µL) were added to a 2 mL HPLC vial. The compounds were partitioned between the layers via vortexing and centrifuged at 1,000 g for 1 min to allow for phase separation. A portion (10 µL) was taken from each layer (autoinjector was set up to draw volume at two different heights) and analyzed by HPLC. The relative concentration in each phase was determined by integrating the area under each peak and comparing the ratio of the areas from the octanol and aqueous layers. A calibration curve was performed to ensure that the concentrations detected were within the linear range of the detector. This procedure was repeated 4 times for each compound.

### Synthesis of Shaker K^+^ channel RNA

A sample of cDNA clone encoding for Shaker voltage-gated K^+^ channel from *D. melanogaster* with inactivation removed^48^ was generously provided by the laboratory of Prof. Francisco Bezanilla at The University of Chicago. The DNA was amplified, linearized with *Not I* enzyme (New England Biolabs, Inc., Ipswich, MA, USA) and transcribed *in vitro* using the T7 promoter mMESSAGE cRNA kit (Ambion, Austin, Tex., USA).

### Expression of Shaker K^+^ channels in Xenopus oocytes

All animal experiments were approved by the University of Guadalajara, CUCEI, Institutional Animal Care and Use Committee. Shaker channel was heterologously expressed in *Xenopus laevis* oocytes. Only mature *Xenopus laevis* frogs (Aquanimals SA de CV, Queretaro, Mexico) were used as oocytes suppliers. A volume of 1-3 mL from the ovary lobes was extracted via survival surgery under anesthesia. Subsequently, oocytes were isolated with the treatment of collagenase type II (Worthington Biochemical Corp., NJ, USA) under mechanical agitation. After the isolation with collagenase, each oocyte was injected with 15-25 ng of RNA encoding for the Shaker ion channel and incubated for 8-12 h at 17 °C in a Standard Oocytes Saline (SOS) solution containing (in mM): 100 NaCl, 1 MgCl_2_, 10 HEPES, 2 KCl and 1.8 CaCl_2_ with 50 μg/mL gentamycin at pH 7.5.

### Recording of K^+^ currents using cut-open voltage clamp

All chemical compounds for this study were acquired from Sigma-Aldrich (Sigma-Aldrich Co., St. Louis, MO, USA) and Chem-Impex International (Chem-Impex International, Inc. Wood Dael, IL, USA) unless otherwise indicated. Electrophysiology measurements were conducted using the methodology of cut-open oocyte voltage clamp (COVC)^49^. For COVC procedures, the internal recording solution contained (in mM): 120 KOH, 2 EGTA, and 20 HEPES. The external recording solution was composed (in mM) by: 12 KOH, 2 Ca(OH)_2_, 105 NMDG (N-methyl-*D*-glucamine)-methylsufonate (MES), and 20 mM HEPES. For measurements carried out at pH of 6.8 and 7.4, the pH of both solutions was adjusted with MES. For measurements carried out at pH = 9.1, HEPES was replaced by 2-(cyclohexylamino)ethanesulfonic acid (CHES).

To quantify the effects upon K^+^ currents of 4AP and 4AP analogs, oocytes that successfully expressed the Shaker channel were voltage-clamped in a COVC station. K^+^ currents were recorded in the same oocyte, first in absence (*I*_*K*_) and then in presence of each drug (*I*_*I*_). *I*_*K*_ was elicited by depolarizing the oocyte membrane with a voltage protocol that consisted in steps of 50 ms from −100 to 50 mV in increments of 10 mV. Then, *I*_*I*_ was achieved by replacing the external solution (top and guard chambers) with a solution containing increasing concentrations of each drug (4AP), 3-fluoro-4-aminopyridine (3F4AP), 3-methyl-4-aminopyridine (3Me4AP), 3-methoxy-4-aminopyridine (3MeO4AP) and 3-trifluoromethyl-4-aminopyridine (3CF_3_4AP). Because 4AP and 4AP analogs block the K_V_ channels in its open conformation^31^, the oocytes were pulsed 5 to 10 times at 10 mV (1 min pulse) until a stable *I*_*I*_ was achieved. The integrity and stability of each oocyte were continuously monitored throughout the experiment.

Ionic currents were amplified and digitized with the Oocyte Clamp Amplifier CA-1A (Dagan Corporation, Minneapolis, MN, USA) and the USB-1604-HS-2AO Multifunction Card (Measurement Computing, Norton, MA, USA), respectively, and controlled with the GpatchMC64 program (Department of Anesthesiology, UCLA, Los Angeles, CA, USA) via a PC. Data were sampled at 100 kHz and filtered at 10 kHz. All the experiments were performed at room temperature (21–23 °C).

### Electrophysiology data analysis

Ion currents recordings were analyzed with Analysis (Department of Anesthesiology, UCLA, Los Angeles, CA, USA) and OriginPro 8 (OriginLab Corporation, Northampton, MA, USA.) programs. The half-maximal inhibitory concentration of 4AP and 4AP analogs (*IC*_*50*_) was determined by fitting the relative current (*I*_*Rel*_ *= I*_*I*_*/I*_*K*_) as a function of the cumulative concentration of each drug ([X]) with the Hill equation:

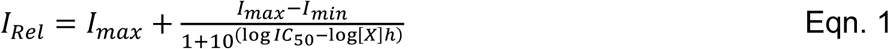

where *I*_*max*_ and *I*_*min*_ are the maximal and minimal value of *I*_*Rel*_, respectively, and *h* is the Hill coefficient, which was typically 1.0 ± 0.1 under our experimental conditions.

The voltage-dependence of blocking by 4AP and 4AP analogs was analyzed in terms of the *IC*_*50*_ as a function of V (*IC*_*50*_*(V)*). *IC*_*50*_*(V)* curves were fitted with a one-step model of inhibition^50,51^, which allowed to determine the fractional distance through the membrane electrical field (δ) that each 4AP analog has to cross to reach its binding site:

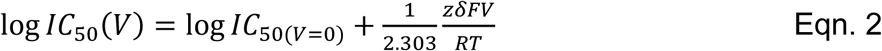

where *IC*_*50(V = 0)*_ is the value of *IC*_*50*_ at V = 0 mV, *F* is the Faraday constant, *R* is the gas constant, *T* is the ambient temperature, and z is the apparent charge. Mean values of data ± standard deviation (s.d.) are given or plotted and the number of experiments is denoted by n. The 95% of confidence interval (IC_95_) is denoted as [Upper limit-Lower limit]; where 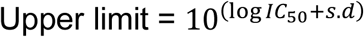 and 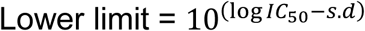.

## Results

### Basicity and lipophilicity of 4AP analogs

The chemical structures and the abbreviations of the 4AP analogs studied are shown in **Figure 2A. Table 1** shows the p*K*_a_ values of these compounds in order of decreasing basicity. As shown on the table: 4AP, 3Me4AP, and 3MeO4AP are more basic with p*K*_a_ values higher than 9, while 3F4AP and 3CF_3_4AP are more acidic with p*K*_a_ values lower than 8. This indicates that the former are mostly protonated at physiological pH, while the latter exist both in the protonated and neutral forms at physiological pH.

**Table 1.**
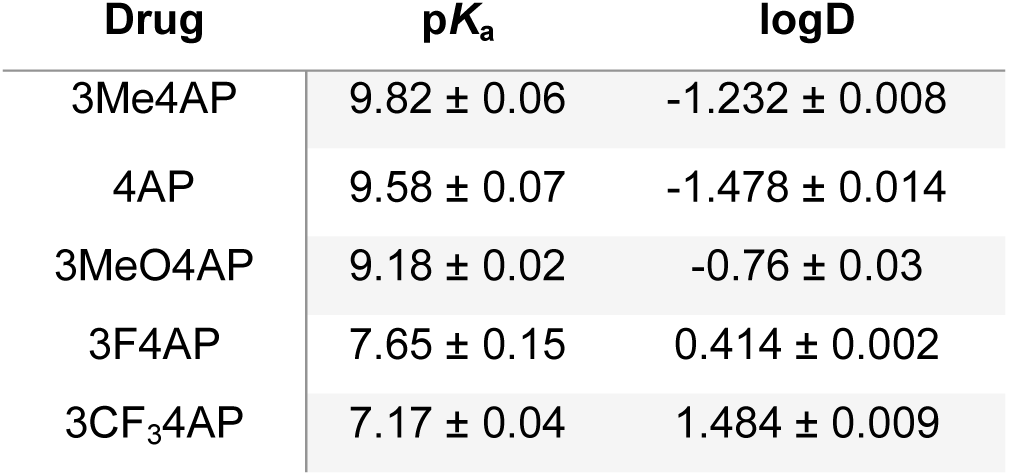
p*K*_a_ and logD (at pH=7.4) values of the 4AP analogs.

In terms of lipophilicity 4AP, 3Me4AP, and 3MeO4AP were found to have octanol/water partition coefficient values at pH 7.4 of −1.48, −1.23 and −0.76 (**Table 1**). This indicates that these compounds preferably partition in the aqueous layer and may have lower penetration of the BBB by passive diffusion. In contrast, 3F4AP and 3CF_3_4AP show partition coefficient values of 0.41 and 1.48 (**Table 1**) indicating that these compounds preferably partition in the octanol layer and may have a faster permeation of the BBB. In fact, 4AP is known to have a slow penetration of the BBB while 3F4AP has a fast BBB penetration.

Both p*K*_a_ and logD trends can be rationalized by the electron-donating or electron-withdrawing nature of the 3-substituent. As the electron-withdrawing strength of the group increases, the dipole of the pyridine nitrogen decreases resulting in a more acidic proton. This lower basicity of the pyridine also results in a greater fraction of the neutral form and higher lipophilicity. A minor deviation from this trend is observed with 3Me4AP. While slightly more basic than 4AP, 3Me4AP is more lipophilic, which can be rationalized by the lipophilic nature of the methyl group compared to a proton in 4AP.

### Blocking capacity of 4AP analogs

K^+^ currents were measured in *Xenopus* oocytes expressing the commonly studied voltage-gated K^+^ channel Shaker from *D. melanogaster*. In order to determine the relative blocking capacity, each drug was applied to the same oocyte at increasing concentrations ranging from 1 to 10,000 μM. Then, the relative current was determined as the ratio between the maximal amplitude of the K^+^ current in the absence and in the presence of each drug. **Figure 2B** shows five representative K^+^ current traces from Shaker elicited at 40 mV, before and after addition of 1,000 μM of each drug. **Figure 2C** shows the relative current as a function of the concentration of each 4AP analog tested. The Hill equation (**Equation 1**) was fitted to the dose-response curves and used to calculate the IC_50_ for each drug at pH 7.4. Hill parameters are summarized in **Figure 2D**. Our results indicate that the relative potency of blocking, from highest to lowest, of these 4AP analogs is: 3Me4AP, 3F4AP, 4AP, 3MeO4AP and 3CF_3_4AP. Specifically, our results show that at pH 7.4 and voltage 40 mV 3Me4AP is approximately 7-fold more potent than 4AP, 3F4AP is comparable to 4AP, and 3MeO4AP and 3CF_3_4AP are approximately 3- and 4-fold less potent than 4AP, respectively. These results are in agreement with the prior observation that small modifications in the 3 position of 4AP are permitted, whereas large modifications significantly diminish the potency of blockage^33^.

### Dependence of the blocking on pH

Since the canonical mechanism describes that only positively-charged molecules can block the channel^41^ and protonation of the drug is dependent on the pH, we studied the blocking of the channel (IC_50_) at pH 6.8, 7.4 and 9.1. To avoid a gradient in pH, both internal and external solutions were replaced.

Panels **A** to **E** of **Figure 3** show the effects on the relative current as a function of concentration for each 4AP derivative at the different pH values. Interestingly, the analogs with high p*K*_a_ (4AP, 3Me4AP and 3MeO4AP) showed higher blocking ability (lower IC_50_) at higher pH, whereas the analogs with lower p*K*_a_ showed higher blocking ability at lower pH. Since 4AP and derivatives bind from the intracellular side^16^, we hypothesize that in the case of the compounds with high p*K*_a_ the limiting factor is the permeability of the drug through the oocyte membrane at low pH. In the case of the compounds with low p*K*_a_, these drugs are able to permeate through the membrane even at low pH and the limiting factor is the fraction of protonated or active form of the drug. **Table 2** summarizes the IC_50_ values of each analog at different pH values calculated by fitting the Hill equation to the data.

**Table 2:** IC_50_ values of 4AP analogs at 0mV: Hill parameters.

| Drug | pH 6.8 |  |  | pH 7.4 |  |  | pH 9.1 |  |  |
| --- | --- | --- | --- | --- | --- | --- | --- | --- | --- |
| | IC<sub>50</sub><br>(μM) | 95% C.I.<br>(μM) | n | IC<sub>50</sub><br>(μM) | 95% C.I.<br>(μM) | n | IC<sub>50</sub><br>(μM) | 95% C.I.<br>(μM) | n |
| 4AP | 210 | 176-244 | 4 | 199 | 190-207 | 6 | 38 | 33-44 | 4 |
| 3F4AP | 82 | 76-88 | 5 | 160 | 140-179 | 5 | 185 | 144-226 | 4 |
| 3Me4AP | 66 | 56-76 | 6 | 39 | 36-42 | 4 | 15 | 13-17 | 5 |
| 3MeO4AP | 357 | 325-390 | 4 | 355 | 327-382 | 4 | 492 | 446-538 | 5 |
| 3CF<sub>3</sub>4AP | 249 | 150-348 | 5 | 690 | 633-747 | 6 | * | * | 4 |

\* Not determined. Relative current vs. concentration did not exhibit a sigmoidal behavior

**Figure 3.**
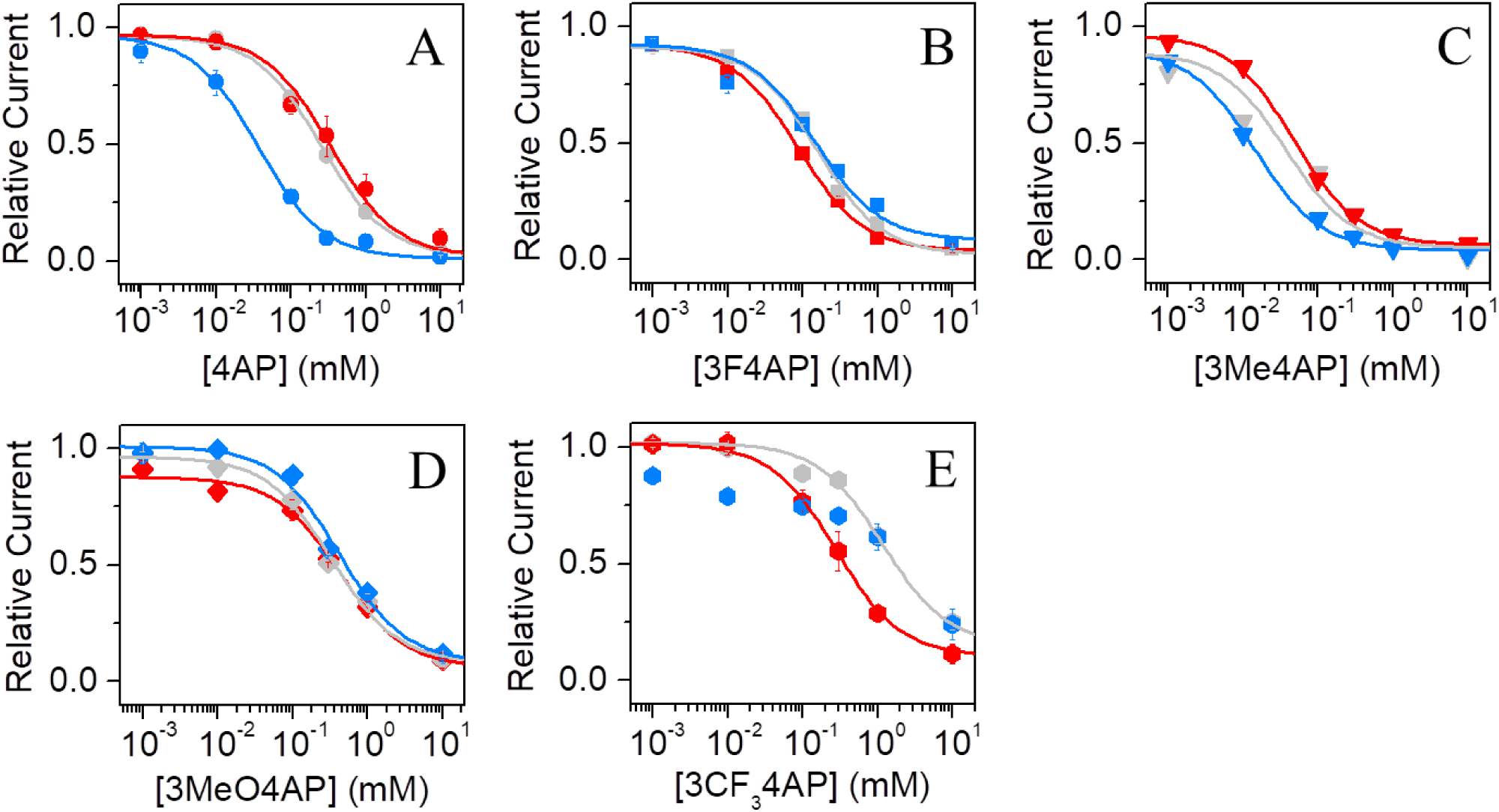
IC_50_ at 0 mV and pH dependence. Relative current *vs.* concentration at 0 mV at pH of 6.8 (red), 7.4 (gray) and 9.1 (blue) of: (**A**), 4AP, (**B**), 3F4AP, (**C**), 3Me4AP, (**D**), 3MeO4AP and, (**E**), 3CF_3_4AP. Continuous lines in panels **A** to **E** represent the fits with the Hill equation (Eqn. 1). Hill parameters are summarized on **Table 2**.

### Dependence of the blocking on voltage

It is known that blocking of K_V_ channels by 4AP involves sequential voltage-dependent rearrangements of the protein^31^. During this process, K_V_ channels must transition to the open conformation before 4AP can bind to its site inside the channel pore. Therefore, it is expected that the blocking of the studied 4AP derivatives will be voltage-dependent as it has been shown for 4AP. For this reason, we studied the IC_50_ at several voltage values. We recorded K^+^ currents from Shaker channel in the range of voltage from −100 to 50 mV (see **Figure S1**) and calculated the IC_50_ of each drug and voltage as described above. Panels **A** to **E** of **Figure 4** show the measured relative current as a function of the concentration of each 4AP analog at several voltage values. For clarity, only representative curves at +10, +30 and +50 mV are shown. From **Figure 4**, it can be observed that the calculated IC_50_ for each 4AP analog increased with voltage (in μM): from 200 to 350 for 4AP, from 160 to 304 for 3F4AP, from 37 to 50 for 3Me4AP, from 310 to 992 for 3MeO4AP, and from 690 to 1150 for 3CF_3_4AP. These results confirm that blocking of the channel is voltage-dependent and that it is more difficult to block the passage of K^+^ ions at higher voltages than at lower voltages.

**Figure 4.**
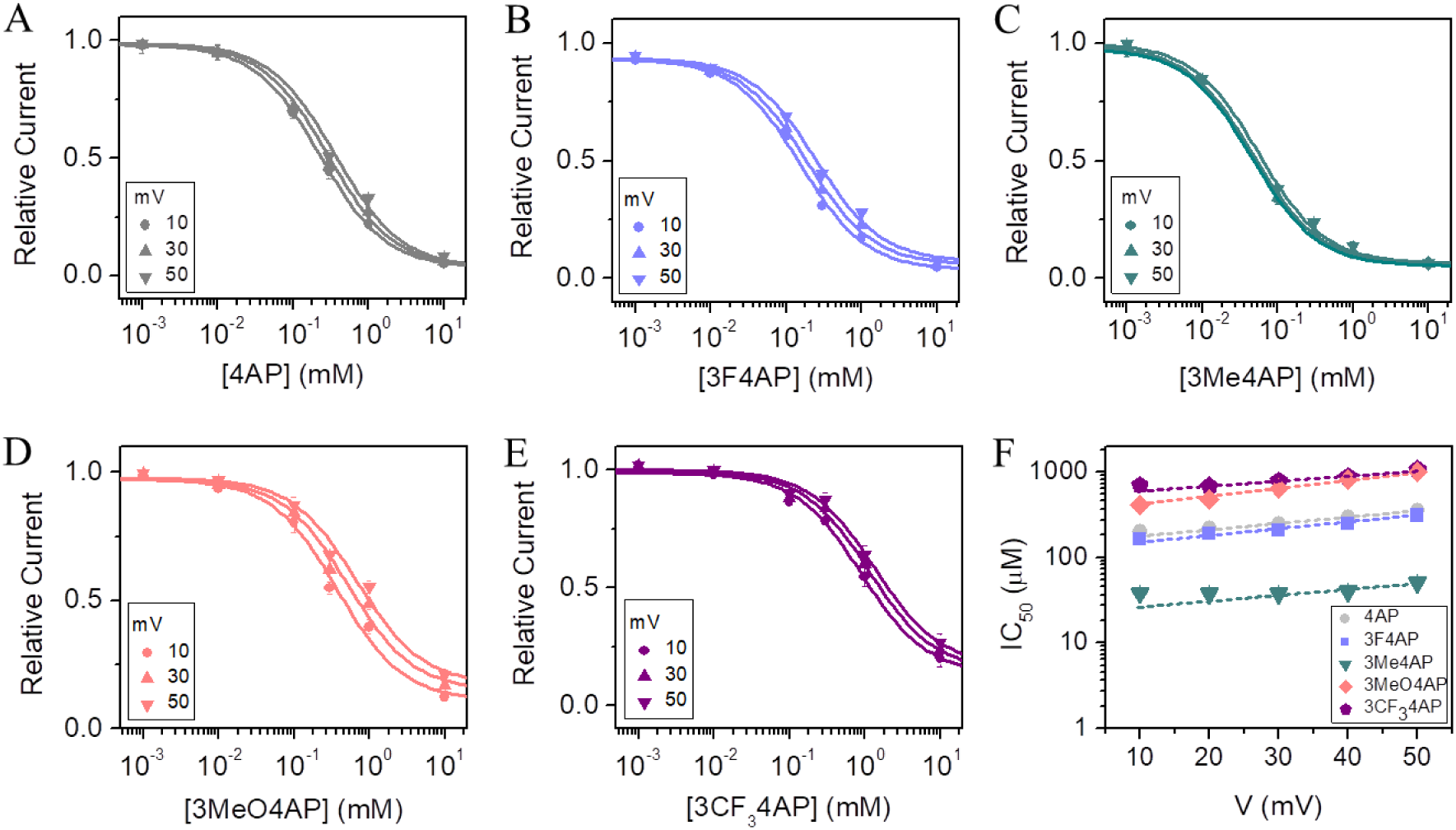
Voltage-dependence of the IC_50_ for each 4AP analog. Relative current as a function of concentration of (**A**) 4AP, (**B**) 3F4AP, (**C**) 3Me4AP, (**D**) 3MeO4AP, and (**E**) 3CF_3_4AP, obtained at different values of voltage. For clarity only the curves obtained at 10, 30 and 50 mV are shown. Solid lines represent the fits of the data with the Hill equation (Eqn. 1). (**F)** IC_50_ *vs.* voltage curves of 4AP analogs determined in the range of voltage from 10 to 50 mV. IC_50_ values were obtained from the analysis of the data of the panels A to E. Dashed lines represent the fits with the Woodhull model (Eqn. 2). Woodhull parameters δ and IC_50_ at V = 0 mV are shown on **Table 3**.

Furthermore, this experiment allowed us to calculate the fractional distance through the membrane electrical field that each molecule travels to bind to its site during the experiment^50^. The Woodhull equation (**Equation 2**) describes that the relationship between IC_50_ and voltage is dependent on δ. By fitting the IC_50_ *vs*. V curves shown in **Figure 4D** to the Woodhull equation, we were able to calculate δ for each 4AP analog. The fitted parameters from this analysis are shown on **Table 3**. Since δ values varied between 0.41 and 0.56, we conclude that these molecules have to travel approximately across one half of the membrane electric field. The similarity of the δ values between 4AP and the other derivatives suggests that these drugs share a common binding site.

**Table 3.** IC_50_ and δ values of 4AP analogs at 0 mV: Woodhull parameters.

| Drug | IC<sub>50</sub> (μM) | $\delta$ | n |
| --- | --- | --- | --- |
| 4AP | 155±12 | 0.41±0.08 | 6 |
| 3F4AP | 122±4 | 0.46±0.10 | 5 |
| 3Me4AP | 21±1 | 0.43±0.10 | 4 |
| 3MeO4AP | 329±17 | 0.56±0.02 | 4 |
| 3CF<sub>3</sub>4AP | 516±51 | 0.46±0.10 | 6 |

\* Mean values are given ± SD

## Discussion

Screening the ability of fluorinated analogs of 4AP to block K^+^ currents in *Xenopus* oocytes expressing Shaker K^+^ channel facilitated the identification of 3F4AP as a potential PET radioligand for demyelination. Subsequent labeling of 3F4AP with ^18^F and PET imaging studies in rodent models of MS confirmed the capacity of this compound to detect demyelinated lesions using PET^33,37^. This compound is in progress to clinical studies and has the potential to advance the imaging of demyelinating diseases.

In PET tracer development, similar to drug development, it is useful to explore multiple derivatives and compare their properties. This process provides confirmation of the target and may result in radioligands with enhanced characteristics. In addition, it is convenient to have ^18^F and ^11^C versions of the same tracer, as each radiolabeling method provides certain advantages that the other lacks. For example, while ^18^F-labeled tracers can be used in sites remote from a cyclotron because of its longer half-life, ^11^C-labeled tracers enable multiple scans (be it more than one ^11^C-scan or a combination of ^11^C/^18^F-scans) on the same subject and the same day.

Here, we studied the efficiency of blocking K^+^ currents by three novel 4AP analogs, namely: 3-methyl-4-aminopyridine (3Me4AP), 3-methoxy-4-aminopyridine (3MeO4AP), 3-(trifluoromethyl)-4-aminopyridine (3CF_3_4AP) (see chemical structures in **Figure 1A**). We selected these compounds as are they are predicted to be permeable to the BBB and are amenable to labeling with ^11^C. During our studies, we found that all of these compounds are able to block Shaker (homolog of mammalian K_v_1.2) channel albeit with different potency. Specifically, 3Me4AP was found to be ∼7 times more potent than 4AP and 3F4AP, whereas 3MeO4AP and 3CF_3_4AP were about 3-4 times less potent. Additionally, we studied the voltage- and pH-dependence of the compounds blocking ability, as these results can provide valuable mechanistic information. From the pH-dependence, we observed that the compounds with lower p*K*_a_ (*i.e.*, 3F4AP and 3CF_3_4AP) are not active at basic pH, presumably because not enough of the protonated/active form is present. At the same time, the compounds with high p*K*_a_ (*i.e.*, 4AP, 3Me4AP and 3MeO4AP) are more active at basic pH as it facilitates membrane permeability. Regarding the voltage-dependence, we found that blocking is more effective at lower voltages than at higher voltages, as the voltage provides a greater driving force for the K^+^ ions. From the voltage-dependence analysis, we were able to estimate using the Woodhull equation that these drugs travel about 50 % of the membrane electrical field in order to bind. This is consistent with prior studies about 4AP binding site^16,29^ and strongly suggests that these drugs share a common binding site.

In summary, we have characterized three novel derivatives of 4AP as potential candidates for therapy and imaging. The physicochemical and pharmacological properties described here will be useful for selecting the compounds with most potential and to explain the differences in terms of kinetics and binding.

## Acknowledgements

J.E.S.R. thanks Dr. Oscar Blanco Alonso, Dr. Jorge Arreola and Dr. Francisco Bezanilla for their support to the Biophysics Laboratory at the University of Guadalajara. We thank Roberto Gastélum Garibaldi for his help with the care and preparation of Xenopus oocytes. We thank Professor Francisco Bezanilla for kindly providing the software for the USB-1604-HS-2AO multifunction card.

## Funding

M.S.R.R. was supported by a fellowship from CONACyT, Mexico (886951), A.D.B. was partially supported by the Faculty Aide Program and the Harvard College Research Program from Harvard University. K.M.R.T. was supported by NIH T32EB013180 (Prof. G. El Fakhri, PI). This work was partially supported by R00EB020075 (P.B.) and PROSNI-UdeG 2017-18 and PRODEP-SEP-2018 (J.E.S.R.).

## Author Contributions

M.S.R.R. performed the COVC experiments and analyzed the data. A.D.B. and K.M.R.T. performed the p*K*_a_ and logD experiments and analyzed the data. P.B. and J.E.S.R. conceived and supervised the project and wrote the manuscript. All authors revised the manuscript.

## Additional Information

The author(s) declare no competing interests.

## REFERENCES

1 Waxman, S. G. & Ritchie, J. M. Molecular dissection of the myelinated axon. Annals of neurology 33, 121–136, doi: 10.1002/ana.410330202 (1993).

2 Trimmer, J. S. & Rhodes, K. J. Localization of voltage-gated ion channels in mammalian brain. Annu Rev Physiol 66, 477–519, doi: 10.1146/annurev.physiol.66.032102.113328 (2004).

3 Jukkola, P. I., Lovett-Racke, A. E., Zamvil, S. S. & Gu, C. K+ channel alterations in the progression of experimental autoimmune encephalomyelitis. Neurobiology of disease 47, 280–293, doi: 10.1016/j.nbd.2012.04.012 (2012).

4 Vacher, H., Mohapatra, D. P. & Trimmer, J. S. Localization and targeting of voltage-dependent ion channels in mammalian central neurons. Physiol Rev 88, 1407–1447, doi: 10.1152/physrev.00002.2008 (2008).

5 Coman, I. et al. Nodal, paranodal and juxtaparanodal axonal proteins during demyelination and remyelination in multiple sclerosis. Brain: a journal of neurology 129, 3186–3195, doi: 10.1093/brain/awl144 (2006).

6 Arroyo, E. J., Sirkowski, E. E., Chitale, R. & Scherer, S. S. Acute demyelination disrupts the molecular organization of peripheral nervous system nodes. The Journal of comparative neurology 479, 424–434, doi:10.1002/cne.20321 (2004).

7 Karimi-Abdolrezaee, S., Eftekharpour, E. & Fehlings, M. G. Temporal and spatial patterns of Kv1.1 and Kv1.2 protein and gene expression in spinal cord white matter after acute and chronic spinal cord injury in rats: implications for axonal pathophysiology after neurotrauma. The European journal of neuroscience 19, 577–589 (2004).

8 Rasband, M. N. et al. Potassium channel distribution, clustering, and function in remyelinating rat axons. The Journal of neuroscience: the official journal of the Society for Neuroscience 18, 36–47 (1998).

9 Wang, H., Allen, M. L., Grigg, J. J., Noebels, J. L. & Tempel, B. L. Hypomyelination alters K+ channel expression in mouse mutants shiverer and Trembler. Neuron 15, 1337–1347 (1995).

10 Sinha, K., Karimi-Abdolrezaee, S., Velumian, A. A. & Fehlings, M. G. Functional changes in genetically dysmyelinated spinal cord axons of shiverer mice: role of juxtaparanodal Kv1 family K+ channels. Journal of neurophysiology 95, 1683–1695, doi: 10.1152/jn.00899.2005 (2006).

11 Fehlings, M. G. & Nashmi, R. Changes in pharmacological sensitivity of the spinal cord to potassium channel blockers following acute spinal cord injury. Brain research 736, 135–145 (1996).

12 Eng, D. L., Gordon, T. R., Kocsis, J. D. & Waxman, S. G. Development of 4-AP and TEA sensitivities in mammalian myelinated nerve fibers. Journal of neurophysiology 60, 2168–2179, doi: 10.1152/jn.1988.60.6.2168 (1988).

13 Kocsis, J. D. Aminopyridine-sensitivity of spinal cord white matter studied in vitro. Experimental brain research. Experimentelle Hirnforschung. Experimentation cerebrale 57, 620–624 (1985).

14 Ritchie, J. M., Rang, H. P. & Pellegrino, R. Sodium and potassium channels in demyelinated and remyelinated mammalian nerve. Nature 294, 257–259 (1981).

15 McCormack, K., Joiner, W. J. & Heinemann, S. H. A characterization of the activating structural rearrangements in voltage-dependent Shaker K+ channels. Neuron 12, 301–315 (1994).

16 Kirsch, G. E., Shieh, C. C., Drewe, J. A., Vener, D. F. & Brown, A. M. Segmental exchanges define 4-aminopyridine binding and the inner mouth of K+ pores. Neuron 11, 503–512 (1993).

17 Choquet, D. & Korn, H. Mechanism of 4-aminopyridine action on voltage-gated potassium channels in lymphocytes. The Journal of general physiology 99, 217–240 (1992).

18 Kirsch, G. E. & Narahashi, T. of action and active form of aminopyridines in squid axon membranes. The Journal of pharmacology and experimental therapeutics 226, 174–179 (1983).

19 Bostock, H., Sears, T. A. & Sherratt, R. M. The effects of 4-aminopyridine and tetraethylammonium ions on normal and demyelinated mammalian nerve fibres. J Physiol 313, 301–315 (1981).

20 Yeh, J. Z., Oxford, G. S., Wu, C. H. & Narahashi, T. Interactions of aminopyridines with potassium channels of squid axon membranes. Biophysical journal 16, 77–81, doi: 10.1016/S0006-3495(76)85663-9 (1976).

21 Gutman, G. A. et al. International Union of Pharmacology. LIII. Nomenclature and molecular relationships of voltage-gated potassium channels. Pharmacological reviews 57, 473–508, doi:10.1124/pr.57.4.10 (2005).

22 Jensen, H., Ravnborg, M., Mamoei, S., Dalgas, U. & Stenager, E. Changes in cognition, arm function and lower body function after slow-release Fampridine treatment. Mult Scler 20, 1872–1880, doi: 10.1177/1352458514533844 (2014).

23 Goodman, A. D. et al. Sustained-release oral fampridine in multiple sclerosis: a randomised, double-blind, controlled trial. Lancet 373, 732–738, doi: 10.1016/S0140-6736(09)60442-6 (2009).

24 Davis, F. A., Stefoski, D. & Rush, J. Orally administered 4-aminopyridine improves clinical signs in multiple sclerosis. Annals of neurology 27, 186–192, doi: 10.1002/ana.410270215 (1990).

25 Stefoski, D., Davis, F. A., Faut, M. & Schauf, C. L. 4-Aminopyridine improves clinical signs in multiple sclerosis. Annals of neurology 21, 71–77, doi: 10.1002/ana.410210113 (1987).

26 Jones, R. E., Heron, J. R., Foster, D. H., Snelgar, R. S. & Mason, R. J. Effects of 4-aminopyridine in patients with multiple sclerosis. Journal of the neurological sciences 60, 353–362 (1983).

27 Hayes, K. C. et al. Preclinical trial of 4-aminopyridine in patients with chronic spinal cord injury. Paraplegia 31, 216–224, doi: 10.1038/sc.1993.40 (1993).

28 Iaci, J. F. et al. Dalfampridine improves sensorimotor function in rats with chronic deficits after middle cerebral artery occlusion. Stroke; a journal of cerebral circulation 44, 1942–1950, doi: 10.1161/STROKEAHA.111.000147 (2013).

29 Caballero, N. A., Melendez, F. J., Nino, A. & Munoz-Caro, C. Molecular docking study of the binding of aminopyridines within the K^+^ channel. Journal of molecular modeling 13, 579–586, doi: 10.1007/s00894-007-0184-9 (2007).

30 Devaux, J., Gola, M., Jacquet, G. & Crest, M. Effects of K+ channel blockers on developing rat myelinated CNS axons: identification of four types of K+ channels. Journal of neurophysiology 87, 1376–1385, doi: 10.1152/jn.00646.2001 (2002).

31 Armstrong, C. M. & Loboda, A. A model for 4-aminopyridine action on K+ channels: similarities to tetraethylammonium ion action. Biophysical journal 81, 895–904, doi: 10.1016/S0006-3495(01)75749-9 (2001).

32 Sherratt, R. M., Bostock, H. & Sears, T. A. Effects of 4-aminopyridine on normal and demyelinated mammalian nerve fibres. Nature 283, 570–572 (1980).

33 Brugarolas, P. et al. Development of a PET radioligand for potassium channels to image CNS demyelination. Sci Rep 8, 607, doi: 10.1038/s41598-017-18747-3 (2018).

34 Basuli, F., Zhang, X., Brugarolas, P., Reich, D. S. & Swenson, R. E. An efficient new method for the synthesis of 3-[^18^F]fluoro-4-aminopyridine via Yamada-Curtius rearrangement. J Labelled Comp Radiopharm 61, 112–117, doi:10.1002/jlcr.3560 (2018).

35 Brugarolas, P., Bhuiyan, M., Kucharski, A. & Freifelder, R. Automated Radiochemical Synthesis of [^18^F]3F4AP: A Novel PET Tracer for Imaging Demyelinating Diseases. J Vis Exp, e55537, doi: 10.3791/55537 (2017).

36 Brugarolas, P., Freifelder, R., Cheng, S. H. & DeJesus, O. Synthesis of meta-substituted [(18)F]3-fluoro-4-aminopyridine via direct radiofluorination of pyridine N-oxides. Chem Commun (Camb) 52, 7150–7152, doi: 10.1039/c6cc02362b (2016).

37 Brugarolas, P., Reich, D. S. & Popko, B. Detecting Demyelination by PET: The Lesion as Imaging Target. Mol Imaging 17, 1536012118785471, doi: 10.1177/1536012118785471 (2018).

38 Berger, S. G., Waser, P. G. & Hofmann, A. Effects of new 4-aminopyridine derivatives on neuromuscular transmission and on smooth muscle contractility. Arzneimittelforschung 39, 762–765 (1989).

39 Sun, W. et al. Novel potassium channel blocker, 4-AP-3-MeOH, inhibits fast potassium channels and restores axonal conduction in injured guinea pig spinal cord white matter. Journal of neurophysiology 103, 469–478, doi: 10.1152/jn.00154.2009 (2010).

40 Kirsch, G. E. & Narahashi, T. 3,4-diaminopyridine. A potent new potassium channel blocker. Biophysical journal 22, 507–512, doi: 10.1016/S0006-3495(78)85503-9 (1978).

41 Howe, J. R. & Ritchie, J. M. On the active form of 4-aminopyridine: block of K+ currents in rabbit Schwann cells. J Physiol 433, 183–205, doi: 10.1113/jphysiol.1991.sp018421 (1991).

42 Lemeignan, M. et al. The ability of 4-aminopyridine and 3, 4-diaminopyridine to cross the blood-brain barrier can account for their difference in toxicity. Advances in the Biosciences 35, 222 (2013).

43 Acorda Therapeutics, I. Vol. Application No.: 02250 (http://www.accessdata.fda.gov/drugsatfda_docs/nda/2010/022250s000TOC.cfm, 2010).

44 Maddison, P. & Newsom-Davis, J. Treatment for Lambert-Eaton myasthenic syndrome. Cochrane Database Syst Rev, CD003279, doi: 10.1002/14651858.CD003279 (2003).

45 Leung, G., Sun, W., Brookes, S., Smith, D. & Shi, R. Potassium channel blocker, 4-aminopyridine-3-methanol, restores axonal conduction in spinal cord of an animal model of multiple sclerosis. Experimental neurology 227, 232–235, doi: 10.1016/j.expneurol.2010.11.004 (2011).

46 Neelamegam, R. et al. in Society of Nuclear Medicine Annual Meeting 486 (2019).

47 Brugarolas, P., Yang, B. Y., Sanchez-Rodriguez, J., Telu, S. & Pike, V. W. in 255th American Chemical Society National Meeting. FLUO 55 (2018).

48 Hoshi, T., Zagotta, W. N. & Aldrich, R. W. Biophysical and molecular mechanisms of Shaker potassium channel inactivation. Science 250, 533–538 (1990).

49 Stefani, E. & Bezanilla, F. Cut-open oocyte voltage-clamp technique. Methods Enzymol 293, 300–318 (1998).

50 Woodhull, A. M. Ionic blockage of sodium channels in nerve. The Journal of general physiology 61, 687–708, doi: 10.1085/jgp.61.6.687 (1973).

51 Hermann, A. & Gorman, A. L. Effects of 4-aminopyridine on potassium currents in a molluscan neuron. The Journal of general physiology 78, 63–86, doi: 10.1085/jgp.78.1.63 (1981).

